# Synthetic genetic circuits that count developmental cues in plant roots

**DOI:** 10.64898/2026.09.28.755210

**Authors:** Soyeon Choi, Jennifer A. N. Brophy

**Affiliations:** Department of Bioengineering, Stanford University, Stanford, CA 94305, USA; Howard Hughes Medical Institute, Stanford University, Stanford, CA 94305, USA

## Abstract

Plants explore soil environments through their roots, which branch repeatedly to access water and soil-borne nutrients. In many plants, root branches are generated by repeated developmental programs and molecular analyses have not identified markers that distinguish primary from later root branches. Consequently, efforts to engineer plant roots for improved water or nutrient acquisition are limited to changes that affect the entire root system. Here, we developed synthetic genetic circuits to count repeating root development cues and enable differential gene expression in successive root branches of the model plant *Arabidopsis thaliana*. The circuits use a root primordium-specific promoter to express large serine recombinases, which irreversibly invert segments of DNA each time a new lateral root is initiated. This converts repeating rounds of root formation into distinct, heritable DNA states. Because root branch specification and outgrowth occur over a ~2 day period, we tested circuit designs for their ability to count long-duration input pulses and developed an architecture that produces distinct fluorescence outputs in primary, secondary, and tertiary roots. By linking repeated developmental cues to distinct genetic outputs, synthetic genetic circuits that count enable root system engineering at the resolution of individual branches and offer a general strategy for dissecting developmental processes that rely on repeated signals.

## Introduction

Root system architecture shapes how plants explore heterogeneous soil environments to acquire water and nutrients (*1, 2*). In some plants, root branches can differ dramatically in size and anatomy, and these differences are associated with specialized roles. For example, rice plants produce both thick and thin lateral roots that either branch repeatedly to expand the root network or specialize in solute uptake (*3, 4*). Whether anatomically indistinguishable root branches can have distinct functions remains unclear.

In *Arabidopsis thaliana*, where root development has been extensively characterized at the molecular level, all root branches are anatomically similar and appear to be generated through repeated deployment of a conserved developmental program. Auxin oscillations at the root tip pattern lateral root stem cells along the length of the primary root, after which these cells divide and differentiate to form secondary roots (*5, 6*). As secondary roots grow, the patterning process repeats to generate higher-order branches, with the same molecular machinery used to create each branch(*7*). While some genes show differences in expression between primary, secondary, and higher-order roots, no promoter has been identified that is exclusively active in a particular branch, limiting our ability to genetically manipulate individual root branches.

Recent advances in synthetic biology have enabled gene expression and gene knockout specifically in lateral roots. Researchers have used the *GATA23* promoter, which is active during lateral root stem cell specification(*8*), to express recombinases or CRISPR-Cas9 components for lateral-root-specific gene expression(*9*) and gene knockout(*10*). These approaches have been used to study lateral root stem cell patterning(*11*) and cell fate plasticity(*12*). However, because *GATA23* is activated during the formation of every lateral root, these tools cannot distinguish secondary from tertiary or higher-order lateral roots. Synthetic genetic circuits that can count stem cell specification events could overcome this limitation by producing different outputs according to the number of times a root has branched. Counting circuits have been demonstrated in *Escherichia coli* using externally applied pulses of arabinose(*13*), but have yet to be developed or applied in multicellular organisms.

Here, we developed a synthetic genetic circuit to count rounds of lateral root initiation and enable differential gene expression in each root branch order. The counting circuit converts transient lateral root initiation events into distinct genetic states using a root-primordium specific promoter and large serine recombinases. We tested circuit architectures based on DNA inversion and deletion to determine how effectively they could distinguish long-duration developmental signals and identified an architecture that produces distinct fluorescence outputs in primary, secondary, and tertiary roots. This genetically encoded counter provides a way to directly test whether anatomically indistinguishable root branches serve distinct functions, and to engineer root system architecture at the level of individual branches. More broadly, the ability to count and differentially respond to repeating signals offers a new approach to dissecting developmental processes that rely on repeated signals.

### Materials and Methods Plasmid construction

DNA constructs were assembled using NEBuilder HiFi DNA Assembly Master Mix (New England Biolabs), based on isothermal DNA assembly (*14*). Fragments were amplified with KAPA HiFi DNA Polymerase (Roche) or Phanta Max Master Mix (Vazyme) and purified with the DNA Clean & Concentrator-5 kit (Zymo Research). PCR comprised initial denaturation at 95–98 °C for 3 min, 10–12 touchdown cycles with a 1 °C decrease in annealing temperature per cycle, and 12–14 cycles at a fixed annealing temperature of 58–66 °C. Extension was performed at 72 °C for approximately 30 s kb−1, followed by a final extension at 72 °C for 2–5 min. Plasmids developed for this study are listed in Supplementary Table S1. Recombinase genes were intronized by insertion of the potato ST-LS1 intron IV2 (*15, 16*) or alpha-tubulin intron 2 into *phiC31, bxb1*, or *cre*, respectively, using NetGene2 splice-site predictions to evaluate the insertions (*17, 18*).

### Bacterial strains and growth conditions

*Escherichia coli* NEB 10-beta (Δ(ara-leu) 7697 araD139 fhuA ΔlacX74 galK16 galE15 e14-ϕ80dlacZΔM15 recA1 relA1 endA1 nupG rpsL (StrR) rph spoT1 Δ(mrr-hsdRMS-mcrBC))(*19*) was used for plasmid cloning and propagation. *Agrobacterium tumefaciens* GV3101 carrying the pSoup helper plasmid was used for plant transformation(*20, 21*). *E. coli* was transformed by heat shock or electroporation, and *A. tumefaciens* by electroporation. Cultures were grown in LB with selective antibiotics for each construct (50 µg/ml Spectinomycin for *E. coli* strains and 10 µg/ml tetracyclin, 50 µg/ml gentamicin, 50 µg/ml spectinomycin and 25 µ/ml rifampicin for *A. tumefaciens* strains) at 250 rpm, at 37 °C for *E. coli* and 28 °C for *A. tumefaciens*.

### Plant material and growth conditions

*Arabidopsis thaliana* Columbia-0 (Col-0) was used as the genetic background for all experiments. All seeds were surface sterilized with 70% ethanol for 1 min and 50% commercial bleach (Clorox, 4.13% sodium hypochlorite) containing 0.05% Triton X-100 (Sigma Aldrich) or Tween20 (Sigma Aldrich) for 5–10 min, then washed and stratified in darkness at 4 °C for 2–3 d before sowing. Soil-grown plants were cultivated in Pro-Mix in a Percival chamber under a 16-h light/8-h dark photoperiod, at 22 °C in the light and 18 °C in the dark, 60% relative humidity, and 70 µmol m−2 s−1 photosynthetic photon flux density. *In vitro*-grown plants were sown on 1X Murashige and Skoog (PhytoTechnology Laboratories) with 1% sucrose (Sigma Aldrich) and 0.6% gelzan (Bioworld) media (pH 5.7) and maintained under a 16-h light/8-h dark photoperiod at 24 °C, 70% relative humidity, and 141 µmol m−2 s−1.

### Generation of transgenic plants

Stable transgenic plants were generated by Agrobacterium-mediated floral dip (*22*). A 2-mL saturated overnight culture initiated from a glycerol stock was diluted 1:1,000 into 50 mL selective broth and grown overnight to reach OD600 of 1–2. Cells were collected at 3,800 xg for 25 min and resuspended in 50–150 mL infiltration solution containing 5% (w/v) sucrose (Sigma Aldrich), 10.5 mM MgCl_2_ (Sigma Aldrich), and 0.03% (v/v) Silwet L-77 (Bioworld) to an OD600 of 0.5–1.0. Inflorescences of Col-0 were dipped in the infiltration solution and dried overnight in dark before moving to a growth chamber. Transgenic seeds were isolated using fluorescence microscopy. All of the plasmids in this study contain the FASTred marker (*23*) for selection of transgenic seeds by screening for red fluorescence. Each T1 transgenic plant was treated as one biological replicate within its construct group.

### Fluorescence imaging and image processing

Widefield images were acquired with a Leica M205 FCA stereomicroscope, Leica K8 camera, with LAS X software version 3.10.2. Fluorescence excitation/emission settings were 436/480 nm for CFP, 500/535 nm for YFP, 470/525 nm for GFP, and 560/630 nm for mCherry, with 1,500-ms exposures and camera gain 1. Reflected-light images were acquired with a 50-ms exposure. Images were acquired with a 1x plan-apochromatic objective at 2.72x zoom. Calibrated pixel sizes were read from image metadata. Tile scans were merged and exported as per-channel 16-bit TIFF files.

Images acquired with the same objective, magnification, exposure, gain, and filter settings were grouped for display. Within each group and channel, the black point was defined as the median of image-specific modal background values, and the white point was the 99.9th percentile of the brightest image. Zero-valued padding pixels from tile stitching were excluded. No gamma adjustment was applied. Channels were pseudocolored and merged additively, with reflected brightfield light weighted at 0.6. Scale bars were calculated from calibrated pixel sizes. Panel-specific display ranges, pixel sizes, and scale-bar lengths are provided as .json files in Supplementary Data S7. Cropping did not alter display ranges.

### Amplicon sequencing and DNA-state quantification

Root tips were harvested for DNA sequencing-based switch characterization, following an approach similar to that used by Greco and colleagues to characterize recombinase-based genetic circuits (*24*). For pSC178 first-transition lines, primary-root tips and three age or length classes of secondary-root tips were harvested from seedlings at 8–10 d after germination. In these lines, LR1 (sec1) denotes the oldest sampled secondary root, LR2 (sec2) denotes an approximately 2-cm-long secondary root, and LR3 (sec3) denotes a 0.5–0.8-cm-long secondary root. These labels describe secondary-root age or length classes and not branch order.

Counter plants were harvested after a parent secondary root had produced at least three tertiary roots longer than 0.5 cm. The three sampled tertiary roots were recorded as ter1, ter2, and ter3. In the counter datasets, tissue labeled “sec_tip” is the tip of the parent secondary root associated with the sampled tertiary roots. Tertiary roots were identified and documented in microscope images before harvest, and the ter1/ter2/ter3 identities were retained to compare fluorescence visibility with the corresponding flipped-read percentage. Each sequencing sample comprised one root segment harvested from one plant; samples were collected from plants across lines. Approximately 0.5 cm of each sampled tissue was excised into sterile eight-strip PCR tubes and mixed with 22 µL Dilution Buffer from the Phire Plant Direct PCR Kit (Thermo Fisher Scientific). Samples were heated at 95 °C for 15 min, and 1–4 µL lysate was used as PCR template. Filtered pipette tips were used throughout to reduce cross-contamination. Amplicon-sequencing primers are listed in Supplementary Table S1. Unpurified PCR amplicons were submitted directly to Quintara Biosciences for AmpPro 4K sequencing.

Analyses were performed on Stanford’s Sherlock computing cluster using the biology environment module followed by the minimap2 (minimap2 2.30-r1287), SAMtools (1.16.1), and python3 modules. Reads were aligned to construct-specific flipped and unflipped reference sequences with minimap2 (*25*), and alignments were converted to BAM format and filtered with SAMtools (*26, 27*). Raw reads were filtered using inclusive length windows of 1,000–1,200 nt for Figure 1 first-transition measurements, 1,753–1,953 nt for Figure 2 first-transition measurements, and 1,383–1,583 nt for second-transition measurements in Figures 2 and 3 and Supplementary Figure S3. Expected PCR products were 1,055 bp for the first-transition assay in Figure 1, 1,853 bp for the first-transition assay in Figure 2, and 1,483 bp for the second-transition assay. Length filtering was intended to retain reads close to the product size while allowing modest variation from partial degradation or sequencing errors. Alignment used the map-ont preset with -Y, --MD, and --cs=long. Secondary and supplementary alignment records were excluded with SAMtools flag filter -F 2304, so only the primary alignments were preserved. Primary alignments with mapping quality 0 were retained to preserve the reads with polymorphisms driven by PCR or sequencing errors. Thus, ambiguous state assignments could remain a potential source of classification bias. Reference FASTAs, assay-specific commands, and the read-counting workflow are provided in Supplementary Data S4. The flipped-read fraction was calculated as n_flipped/(n_flipped + n_unflipped) for each tissue and developmental transitions (first and second flipping by the branching cue). Version 3 circuits delete the first-transition assay substrate during intended progression, so percentages calculated from residual first-transition molecules would be biased and were not interpreted quantitatively. Sequencing in counters were done to compare fluorescence visibility with DNA-level flipping in selected tissues, not to rank counter functionality across versions.

**Figure 1.**
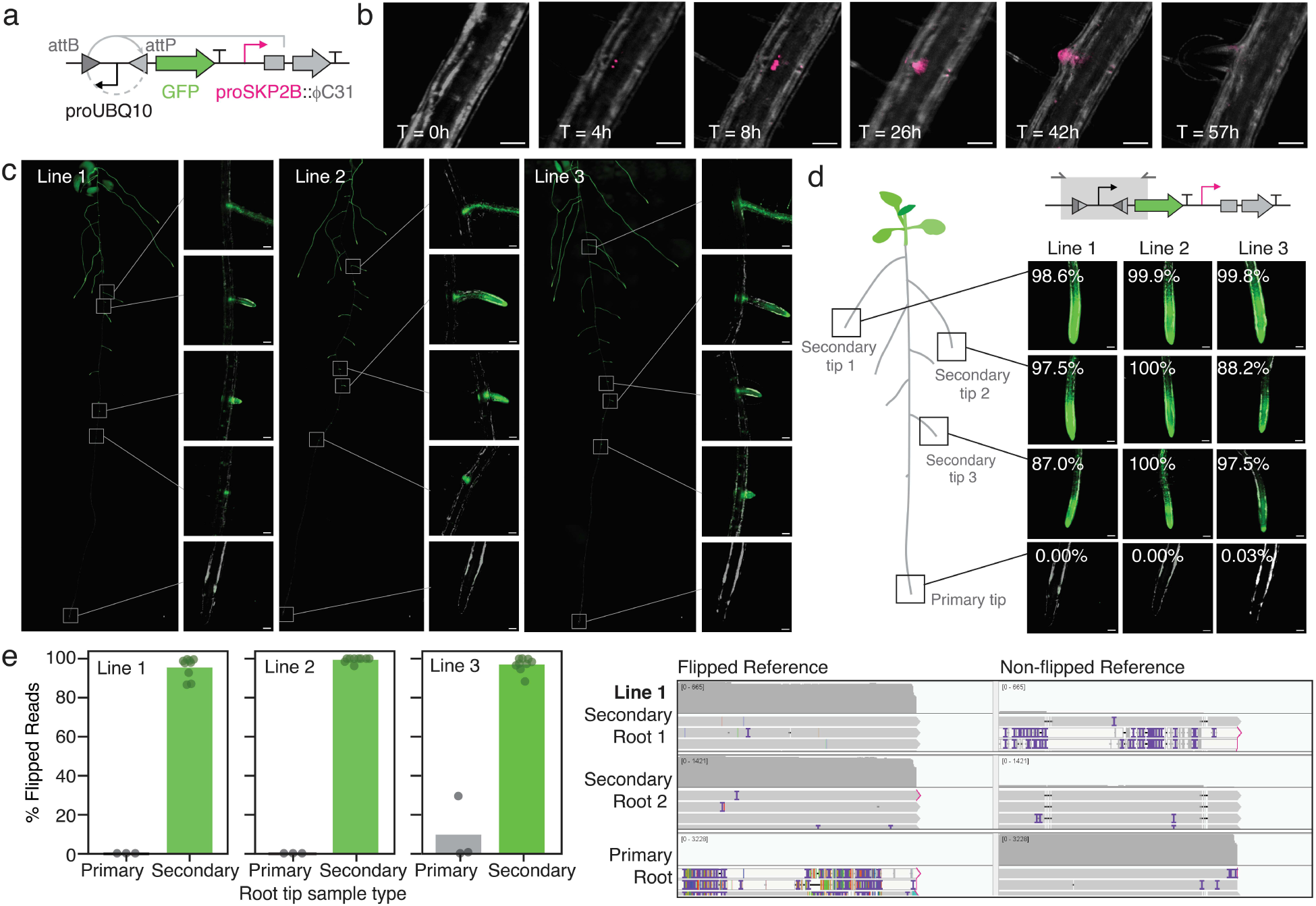
A genetic switch distinguishes secondary roots from the primary root. (a) Schematic of the recombinase-based switch designed to convert transient *SKP2B[0*.*5kb]* promoter activity into a stable output. PhiC31 expression from the *SKP2B[0*.*5kb]* promoter inverts ~0.7kb long DNA segment containing a *UBIQUITIN 10* promoter to initiate expression of GFP. (b) Time-lapse imaging of a proSKP2B[0.5kb]::mGFP6 reporter shows transient reporter activity during lateral root primordia development that declines after root emergence. Scale bars, 100µm. (c) Fluorescence images of T2 plants carrying the switch from (b) in three independent transgenic lines (lines 1-3). Images and insets overlay GFP and brightfield channels. Scale bars, 100µm. (d) Amplicon sequencing of root tips from the same three lines. Half centimeter samples of the primary root, oldest secondary root (secondary tip 1), a ~2-cm-long secondary root (secondary tip 2), and a 0.5–0.8-cm-long secondary root (secondary tip 3) were collected for imaging, DNA extraction, PCR amplification, and sequencing. PCR amplicon reads were mapped to flipped and non-flipped reference sequences to determine flip frequency. Insets show the percent of reads that aligned to the flipped sequence. Below, representative sequencing tracks show reads from line 1 mapped to flipped and non-flipped reference sequences. (e) Activation of the genetic switch in primary and secondary root tips measured by amplicon sequencing for three T2 plants per transgenic line (n = 9 secondary root tip and n = 3 primary root tip samples per line).

**Figure 2.**
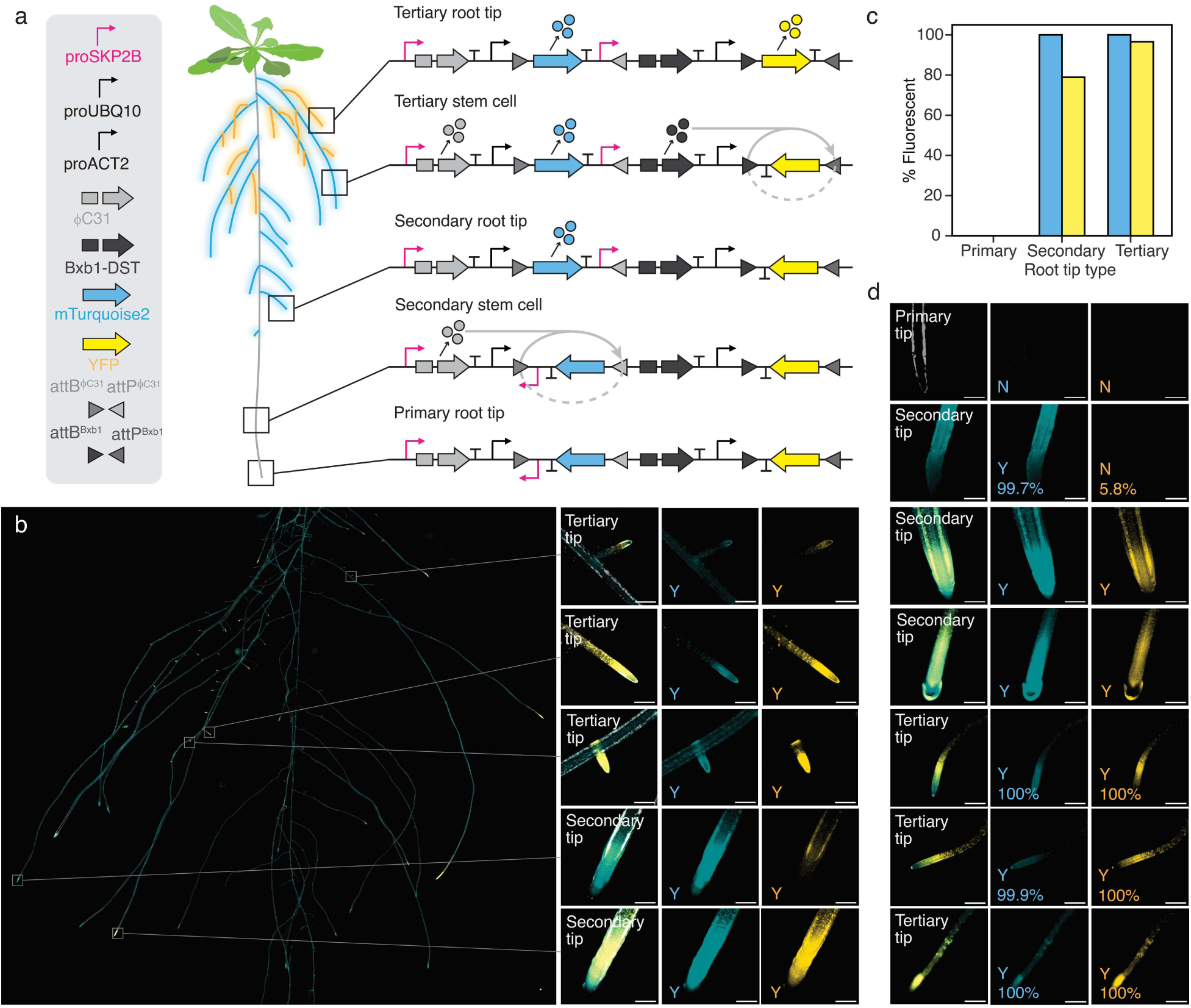
Initial synthetic genetic circuit makes double counting errors. (a) Schematic of the synthetic genetic circuit designed to count root branch initiation events (version 1). Plants carrying the counter should express no fluorescent protein in the primary root, CFP exclusively in lateral roots and higher, and YFP in tertiary roots and higher. Schematic shows the expected DNA state at each stage of primary, secondary and tertiary root development. (b) Images of a representative T1 plant carrying the counter from (a) (Line 2) in independent or merged CFP (cyan), YFP (yellow) and reflected light (grey) channels. Insets show secondary and tertiary root tip signal with Y and N indicating root tips scored as either fluorescence-positive (Y) or fluorescence-negative (N) for CFP and YFP. (c) Percentage of primary, secondary and tertiary root tips with detectable CFP or YFP in the plant shown in (b). One primary, 19 secondary, and 89 tertiary root tips were scored. (d) Representative root tip images from transgenic plants carrying the version 1 counter shown in (a) (Line 2). Y and N indicate root tips scored as either fluorescence-positive (Y) or fluorescence-negative (N) for CFP and YFP. Percentages show the fraction of amplicon sequencing reads that align to flipped sequence marking completion of the first count (CFP output) or second count (YFP output) in the pictured root tip. Additional T1 plants and their corresponding analyses are shown in Supplementary Figure S2. Scale bars, 250µm.

**Figure 3.**
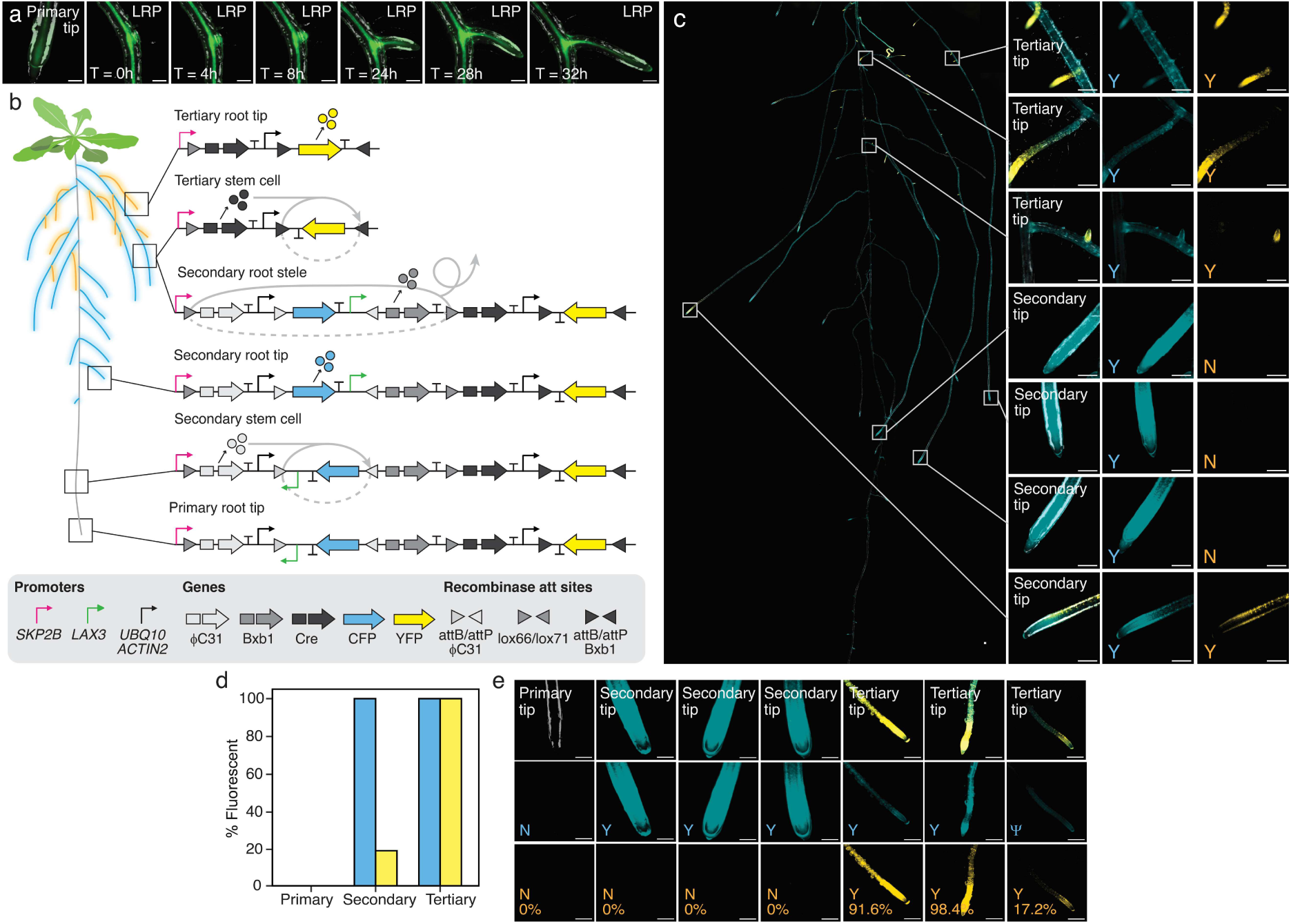
State-gated deletion improves counter performance. (a) Time-lapse imaging of a proLAX3::mScarlet3 reporter shows expression in newly expanding and mature stele tissues. Scale bars, 100µm. (b) Schematic of the synthetic genetic circuit designed to count root branch initiation events (version 3.2) and the expected DNA states during root development. Plants carrying the counter should express no fluorescent protein in the primary root, CFP exclusively in secondary roots, and YFP in tertiary roots. (c) Images of a representative T1 plant carrying the counter from (b) (Line 1) in CFP, YFP, and merged channels. Insets show secondary and tertiary root tip signal with Y and N indicating root tips scored as either fluorescence-positive (Y) or fluorescence-negative (N) for CFP and YFP. Scale bars, 250µm. (d) Percentage of primary, secondary and tertiary root tips with detectable CFP or YFP in the plant shown in (c). One primary, 26 secondary, and 55 tertiary root tips were scored. (e) Selected root tip images from transgenic plants carrying the version 3.2 counter shown in (a) (Line 1). Y and N indicate fluorescence detection in CFP or YFP channels. Percentages show the fraction of amplicon sequencing reads that align to the second inversion (YFP) in the pictured root tip. Additional T1 plants and their corresponding analyses are shown in Supplementary Figure S5. Scale bars, 250µm.

### Data visualization and statistical reporting

Fluorescence incidence and DNA-state fractions across different versions of counters were analyzed as separate outcomes. For visual scoring, image-display brightness was increased after acquisition to aid detection of weak fluorescence. Roots or root tips with any detectable CFP or YFP signal were counted as positive for the corresponding reporter. Fluorescence incidence was the percentage of scored roots or root tips with detectable CFP or YFP, not the percentage of fluorescent cells within a tissue.

Fluorescence analyses compared counter version-specific groups of independent T1 transgenic plants [version 1 (v1), n = 7; version 3.1 (v3.1), n = 6; and version 3.2 (v3.2), n = 9]. Each event contributed one percentage for each combination of root order (primary, secondary, and tertiary) and reporter (CFP and YFP). The combination of counter version and Line identified an event and root-order measurements were paired within an event. Percentages were calculated as 100 X positive roots or root tips divided by the total scored for each event, root order, and reporter, without rounding before analysis. One primary root or root tip was scored per T1 plant. Signal-Positive and total counts are provided in Supplementary Table S2. Independent T1 events were weighted equally within each construct group; roots within a plant were not treated as independent biological replicates. We report medians, interquartile ranges, ranges, and individual-event observations.

Between-version differences in secondary- and tertiary-root YFP incidence were assessed using two-sided exact Wilcoxon–Mann–Whitney rank-sum tests with ties retained (Supplementary Fig. S6a). These tests compare independent counter versions without assuming normally distributed plant-level percentages and thus ‘exact’ calculations accommodate the small sample sizes and tied observations. All three version pairings were tested for each root order, with Holm correction across the six comparisons to control the family-wise error rate (*28*). Exact p values were calculated from all group-label allocations using the absolute deviation of the rank sum from its null mean.

Within each version (Supplementary Fig. S6b), YFP incidence across root orders was compared using two-sided exact paired sign tests, with Holm correction across the nine comparisons. Pairing accounts for measurements from the same plant; sign tests assess the direction of within-plant differences without requiring their magnitudes to follow a normal or symmetric distribution. Zero differences were excluded from each sign test; an all-zero contrast was assigned p = 1. Constant outcomes were summarized descriptively. Adjusted p < 0.05 was the reporting threshold. Nonsignificant results do not necessarily establish equivalence, and these comparisons do not isolate causal effects of individual circuit components.

Statistical analyses and plots were generated in R 4.6.0 (*29*), with ggplot2 4.0.3 used for plotting (*30*). A single R is provided in Supplementary Data S5.

## Results

### Engineering a lateral root-specific genetic switch as a building block for counting

To establish a foundation for genetic circuits that count root branches, we first built a switch that converts transient lateral root initiation signals into permanent genetic memory. This switch uses a lateral root primordia-specific promoter to express the large serine recombinase ϕC31, which inverts a ~0.7kb segment of DNA encoding an Arabidopsis *UBIQUITIN 10* promoter (*21*) to activate expression of GFP (Fig. 1a). This DNA inversion is irreversible and GFP expression should be maintained in all cells derived from the lateral root primordium. Since the canonical marker for lateral root founder cells, *GATA23*, has been reported to be leaky(*8, 9*), causing substantial recombinase-mediated DNA inversions in the primary root when used to generate similar switches, we selected a 0.5kb fragment of the promoter for *S-PHASE KINASE-ASSOCIATED PROTEIN 2B* (proSKP2B[0.5kb]) (*31*) as the basis for our switch. In alignment with prior studies, a proSKP2B[0.5kb]::GFP reporter showed specific activity in lateral root primordia with GFP expression first appearing at stage I of lateral root development, when founder cells undergo their initial asymmetric division, and persisting for approximately 44–48 hours until the lateral root fully emerged (Fig. 1b).

We assessed switch performance in transgenic plants using two complementary approaches. First, using T2 plants derived from three independent transformation events (Lines 1-3), we scored the percentage of secondary roots with visible GFP signal and found that all secondary roots were fluorescent, which is consistent with robust expression of ϕC31 in lateral root primordia (Fig. 1c). Then, we harvested the terminal 0.5cm of the primary root and three secondary roots from each plant, extracted DNA, and amplified the region spanning the flipped segment by PCR to assess switch efficiency through sequencing, as recently described (Fig. 1d). Amplicons were sequenced by nanopore long read sequencing and reads were aligned to flipped and non-flipped references sequences. Amplicon sequencing confirmed near-100% flipping in secondary roots and no detectable flipping in most primary roots. In total, 27 secondary root tips were sequenced with an average flipping efficiency of 95-99% across independent lines (Fig. 1e). In contrast, eight of the nine primary root samples had 0-0.03% flipped reads, with the remaining relatively low-quality sample showing 29.4% flipping. Together, these measurements demonstrate near-complete flipping in secondary roots with minimal primary root activity, establishing a persistent memory module for the counter.

### Implementation of a synthetic genetic circuit that counts in plant roots

To convert repeated rounds of lateral root formation into multiple, distinct genetic states, we designed a two-step recombinase counter modeled on the DNA recombinase-based counter first demonstrated in *Escherichia coli (13)*. In this *E. coli* counter, a cascade of recombinases invert nested DNA segments to count the number of externally applied arabinose input pulses (*13*). In our design, the *SKP2B[0*.*5kb]* promoter is used to express recombinases that generate a stepwise sequence of DNA states corresponding to root branch order (Fig. 2a). The first round of *SKP2B[0*.*5kb]* promoter activation leads to expression of ϕC31 and inversion of a segment containing a second *SKP2B[0*.*5kb]* promoter and the blue fluorescent protein, mTurquoise2.

This inversion activates expression of mTurquoise2 in secondary roots and aligns the second *SKP2B[0*.*5kb]* promoter with Bxb1, another large serine recombinase. We tagged Bxb1 with SAUR-derived RNA-destabilization element DST (*9*) to shorten Bxb1 mRNA persistence and reduce potential for premature recombinase activity (Fig. 1b). A second round of *SKP2B* activation leads to expression of Bxb1 and inversion of a second DNA segment containing yellow fluorescent protein, YFP. This inversion activates expression of YFP in tertiary root primordia and the resulting tertiary roots. Plants carrying this circuit should therefore express no fluorescence in the primary root, mTurquoise2 in secondary roots and above, and YFP in tertiary roots and above.

We generated stable transgenic *Arabidopsis* lines carrying this version 1 counter and imaged whole root systems to assess whether fluorescent output tracked root branch order as designed (Fig. 2b, Supplementary Fig. 2). In a representative plant (Line 2), CFP was detected in nearly all secondary and tertiary root tips, consistent with the first round of *SKP2B*-driven recombination occurring reliably during secondary root initiation (Fig. 2b, c). However, YFP, which should only appear following the second round of recombination in tertiary root primordia, was also detected in a substantial fraction of secondary root tips (Fig. 2b, c). Of 19 secondary root tips scored in this plant, 78.9% showed detectable YFP signal. Thus, this circuit is prone to double counting errors in which both rounds of DNA inversion occur during development of the first lateral root primordium. Amplicon sequencing of individual root tips confirmed double counting errors. Representative secondary root tips contained a high percentage of reads mapping to the second flip (YFP-expressing state) (Fig. 2d). This result indicates that the version 1 counter is prone to double counting and necessitates optimization beyond DST-mediated destabilization of Bxb1 mRNA.

### Deletion-gated architectures suppress premature output and enrich YFP in tertiary roots

The double counting observed in the version 1 circuit likely results from a mismatch in the kinetics of root development and recombination. Friedland et al.’s original DNA recombinase-based counter was designed to respond to brief pulses of an external inducer and was unable to count pulses longer than 10 hours(*13*). In contrast, *SKP2B[0*.*5kb]* promoter activity persists for approximately 44–48 hours during lateral root primordium development (Fig. 1b), likely allowing both recombination events to occur within the same developmental window. To prevent double counting, we designed a second-generation circuit that introduces an obligatory delay between the two counting events using a third recombinase (Cre)(*32*) expressed from the *LIKE AUXIN RESISTANT 3 (LAX3)* promoter(*33*), which is active in mature stele (Fig. 3a). In this design, Cre-mediated deletion must occur before Bxb1 can be expressed. Because *LAX3* activity begins only after a secondary root has matured beyond its *SKP2B* pulse, *Bxb1* can only be expressed during a subsequent round of lateral root initiation, which prevents the first pulse from causing a second count.

We generated stable transgenic *Arabidopsis* lines carrying two versions of this deletion-gated counter and imaged whole root systems to assess whether fluorescent output tracked root branch order as designed (Fig. 3b, Supplementary Figs. 3-5). The first version (3.1) tagged Bxb1 with the SAUR-derived DST RNA-destabilization element used in the plant integrase toolbox (*9*) and our version1 counter to further limit Bxb1 availability (Supplementary Fig. 3a). The second version (3.2) retained the corresponding deletion-gated design without DST to test whether counting could be achieved without additional attenuation (Fig. 3b).

Of the two deletion-gated designs, version 3.2, lacking the DST destabilization element, produced substantially better counting accuracy than version 3.1. In a representative 3.2 T1 plant (Line 1), CFP was detected in nearly all secondary and tertiary root tips, consistent with the first round of *SKP2B*-driven recombination occurring reliably during secondary root initiation (Fig. 3c, d). Of 26 secondary root tips scored in this plant, only 19.2% showed detectable YFP signal, a substantial reduction from the ~80% seen in the version 1 circuit, and 100% of the 55 tertiary root tips were positive for both CFP and YFP. Because Cre-mediated deletion removes the sequence needed to detect the first inversion, amplicon sequencing can only measure completion of the second count. Nevertheless, amplicon sequencing of individual root tips confirmed version 3.2 counting. Representative tertiary root tips showed high fractions of reads aligned to the second inversion (91.6-98.4% in two representative tips) (Fig. 3e). Version 3.2 also produced YFP-positive tertiary roots in all nine T1 plants measured for this study, with a median incidence of 73.3% (range, 20–100%) (Supplementary Fig. 5). Together, these results establish version 3.2 as an effective deletion-gated counter that converts successive rounds of lateral root initiation into differential labeling of secondary and tertiary roots.

In comparison, version 3.1 carrying the DST-destabilized Bxb1, showed minimal YFP expression even in tertiary root tips (Supplementary Figs. 3-4). This suggests that the DST element attenuated Bxb1 expression beyond the threshold needed for efficient recombination. In a representative 3.1 T1 plant (Line 1), CFP was detected in nearly all secondary and tertiary root tips, consistent with the first round of *SKP2B*-driven recombination occurring reliably during secondary root initiation (Supplementary Fig. S3b-d), but none of the secondary root tips scored in this plant showed detectable YFP signal. Thus, the DST tag makes this design more stringent, however only 29.0% of the 62 tertiary root tips were positive for both CFP and YFP. Across all six independent lines, four produced YFP-positive tertiary roots and median tertiary-root incidence was 18.1% (range, 0–65.4%). Thus, the version 3.1 counter underperformed relative to version 3.2 and DST-based attenuation of Bxb1 expression was unnecessary for robust counting *in planta*.

## Discussion

Repeated signals, where the same molecular machinery is used repeatedly to generate distinct outcomes, is common in development. In these cases, conventional genetic tools can control responses to the presence of a signal, but cannot distinguish between successive occurrences of that signal. A genetic counter provides access to otherwise inaccessible points along a developmental trajectory by making gene expression dependent on developmental history rather than signal presence alone. This strategy could be extended to other repeating developmental cues, such as the recurring hormonal signals that regulate axillary bud activation during shoot branching, or used to trigger a response after a defined number of environmental signals. In this way, counting could provide a means to encode environmental history into gene expression and manipulate developmental responses based on the cumulative experience of a plant rather than its response to a single stimulus.

In the Arabidopsis root system, the ability to count lateral root stem cell specification events provides new opportunities to study the functional contributions of individual branches and to engineer root architecture. For example, selectively removing root hairs from one branch type while leaving them intact on others could reveal how individual branches contribute to whole-plant fitness. The same approach could be used to control other branch-specific traits, such as gravitropic responses, allowing different branches to adopt distinct growth trajectories based on their developmental history. Such control could ultimately enable user-defined root system architectures in which different branches are programmed to occupy distinct regions of the soil.

More broadly, the ability to couple gene expression to the number of times a developmental event has occurred could provide a framework for controlling developmental processes in other multicellular organisms. For example, a genetic counter that differentially responds to successive Vascular endothelial growth factor (VEGF) signaling events could be used to alter endothelial cell behavior after a defined number of signaling events, potentially allowing vascular development to be manipulated according to developmental history rather than VEGF signaling alone (*34*). Similarly, counting successive Notch-dependent cell fate decisions could enable different gene expression programs to be activated at distinct stages of iterative cell specification, providing a means to manipulate cell fate based on the number of prior signaling events (*35*). Thus, genetic counters could add developmental history as a new dimension of genetic control.

## Acknowledgements

We thank Dr. Jon Cody, Dr. Colby Starker and Dr. Dan Voytas at the University of Minnesota for providing the Cre recombinase used in this study.

## Author contributions

S.C. and J.A.N.B. designed the study. S.C. conducted experiments, curated and analyzed, visualized and interpreted data, and drafted and edited the manuscript. J.A.N.B. supervised the study, data visualization and interpretation, and edited the manuscript.

## Data, code, and materials availability

Sequencing data, analysis and plotting code, and image-processing presets are described in Supplementary Data S1–S7. Supplementary Data S4 contains figure-specific amplicon-analysis scripts and alignment references. Supplementary Data S5 contains the R script for statistical analysis and plotting. Fluorescence incidence measurements, statistical analyses and amplicon sequencing results are reported in Supplementary Table S2-4. Plasmids will be available through Addgene.

