## Supplementary Figures for "Synthetic genetic circuits that count developmental cues in plant roots"

**This PDF file includes:**

Supplementary Figures S1 to S6
Captions for Supplementary Tables S1 to S7

**Other Supplementary Materials for this manuscript include the following:**

Supplementary Data S1 to S7
Supplementary References

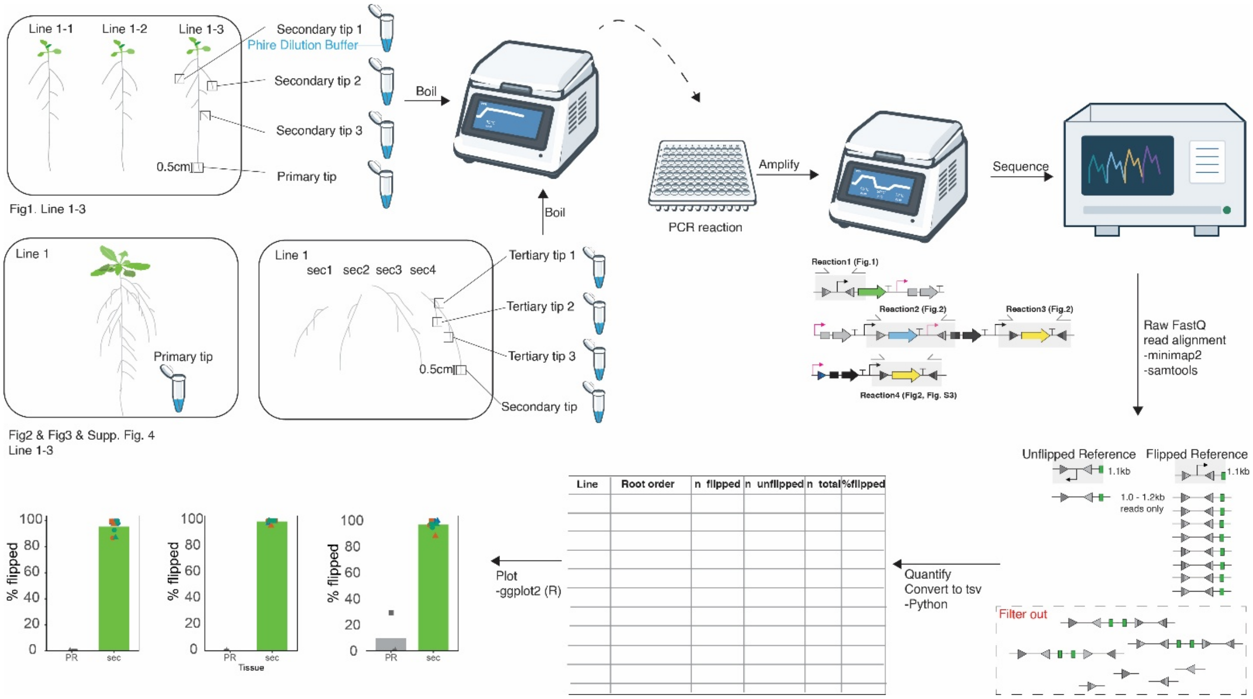


**Supplementary Figure S1. Sampling and sequencing workflow for DNA-state measurements.** Root tip sampling strategy. For switch experiments (Figure 1), the primary root tip and three secondary root tips were sampled from each plant. Secondary roots were selected as follows: the oldest (most shootward) secondary root, an ~2cm long secondary root, and the youngest (most rootward) secondary root that was at least 0.5cm long. For counter experiments (Figures 2-3), the primary root tip, three secondary root tips, and three tertiary root per secondary were sampled from each plant. The terminal 0.5cm of each root was collected for analysis. DNA was extracted from each root tip sample by placing the tissue in Phire Dilution Buffer and heating to 95 °C for 15 min.

One or two PCR reactions were run using DNA extracted from each root tip sample. Reaction 1 was run to measure DNA flipping in the switch construct (Figure 1). Reactions 2-3 measure the first and second DNA flips (counts) in the version 1 counter (Figure 2). Reaction 4 measures the second DNA flip in the version 3.2 counter (Figure 3). PCR products were sequenced using primordium premium PCR (nanopore) sequencing. FASTQ reads were filtered by length to remove aberrant reads and aligned to construct-specific unflipped and flipped reference sequences with minimap2. The dashed box illustrates reads excluded by length filtering.

Flipped read percentages were calculated as 100 × n_flipped/(n_flipped + n_unflipped). Primer sequences, sample-level DNA-state measurements, and amplicon sequence analysis scripts are provided in Supp. Table S1, Supp. Table S4, and Supp. Data S4, respectively.

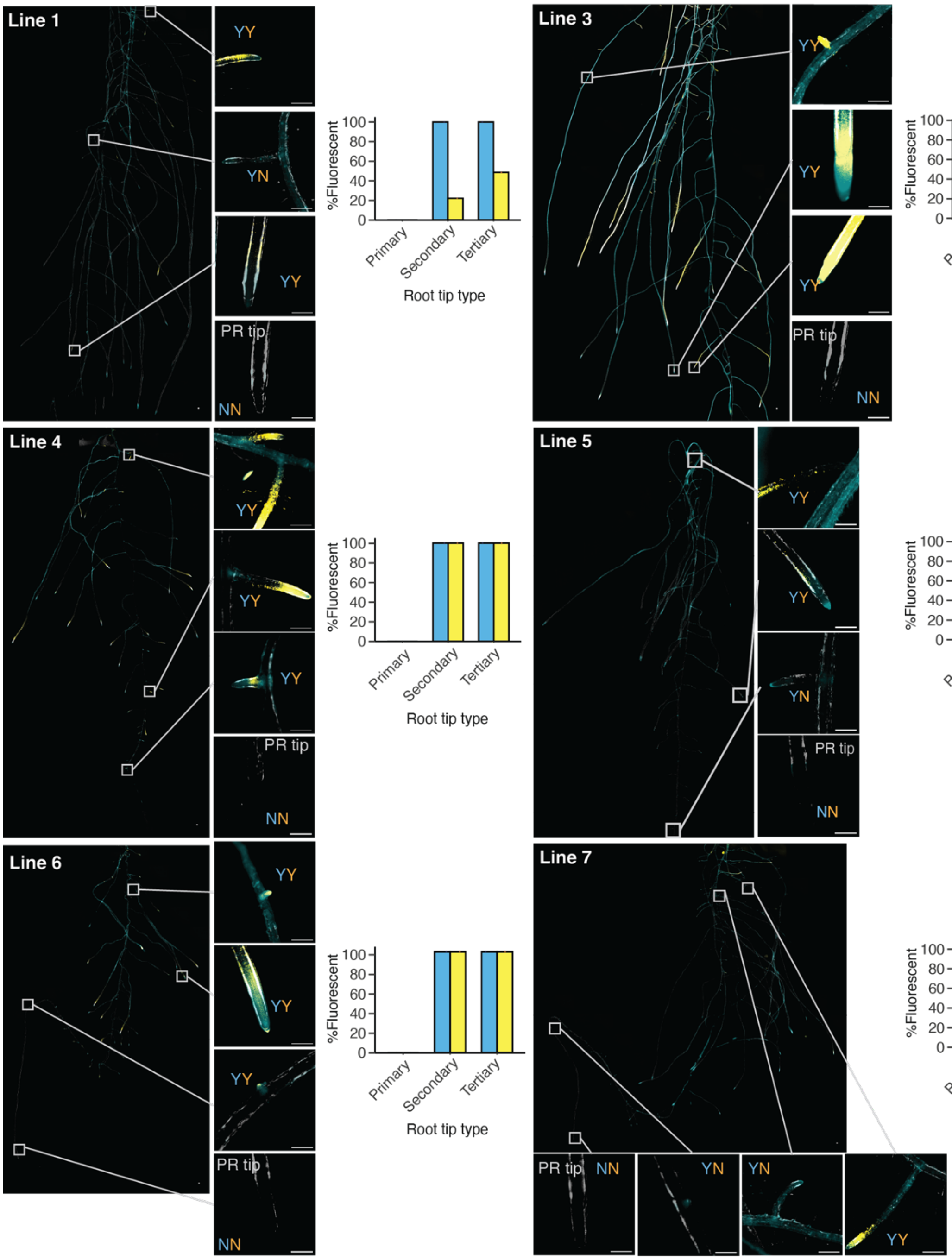


**Supplementary Figure S2. Additional T1 transgenic lines for version 1 counter.** Root-system fluorescence images and percentage of primary, secondary and tertiary root tips with detectable CFP or YFP signal for six additional T1 plants (Lines 1 and 3–7) carrying the version 1 counter (pSC235); Line 2 is shown in Figure 2. Images show merged CFP (cyan), YFP (yellow), and reflected-light (gray) channels. Insets show secondary and tertiary root tip signal with Y and N indicating root tips scored as either fluorescence-positive (Y) or fluorescence-negative (N) for CFP (blue) and YFP (yellow). Each plot shows percentage data for one T1 plant. Signal-positive and total counts are provided in Supplementary Table S2, and across-plant statistical comparisons are reported in Supplementary Table S3. Scale bars, 250 µm.

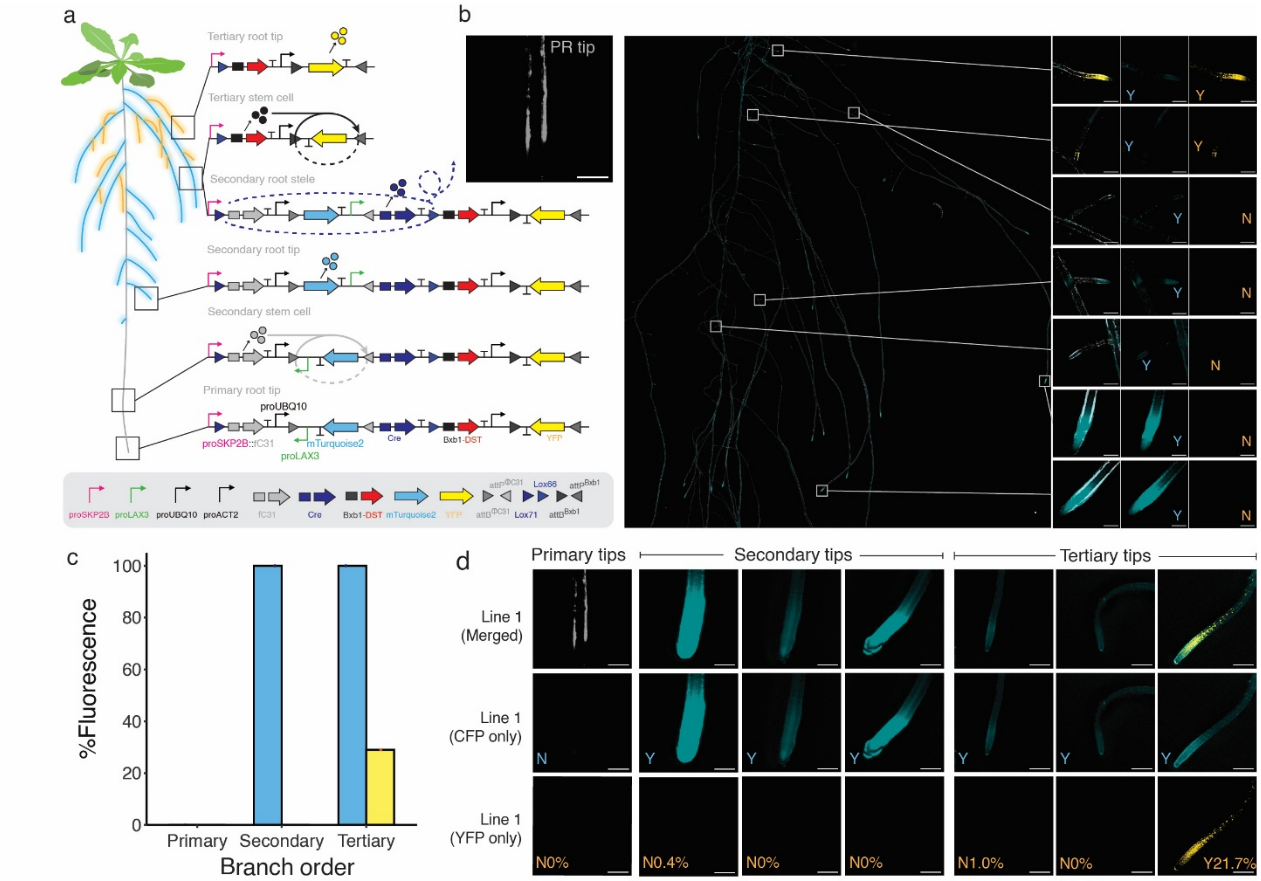


**Supplementary Figure S3. Schematic and performance of state-gated deletion counter with DST-tagged Bxb1.** (a) Schematic of the counter architecture and intended progression using ϕC31 for the first inversion, Cre/lox71/lox66 for deletion, and Bxb1-DST for the second inversion.

**(b)** Representative Line 1 whole-root fluorescence, primary-root-tip inset, and enlarged regions displayed as merged channels, CFP, and YFP.

**(c)** Event-specific fluorescence incidence by root order. For the representative v3.1 T1-1 plant, 18 of 62 tertiary roots were YFP-positive (29.0%). Panel c comprises one primary root or tip, 11 secondary roots or tips, and 62 tertiary roots or tips. All signal-positive counts are provided in Supplementary Table S2.

**(d)** Selected primary, secondary, and tertiary root-tip images from Line 1. Y and N denote fluorescence detection while percentages in the YFP row indicate classified second-transition reads in the corresponding tissue. Retained CFP is a reporter observation and does not measure deletion completion. Additional events and their corresponding plots are in Supplementary Figure S4. Data and analyses are in Supplementary Tables S2–S4. Each sequencing annotation corresponds to one harvested root segment and permits a sample-level comparison with fluorescence visibility, not a direct comparison of overall counter functionality. Scale bars = 250μm.

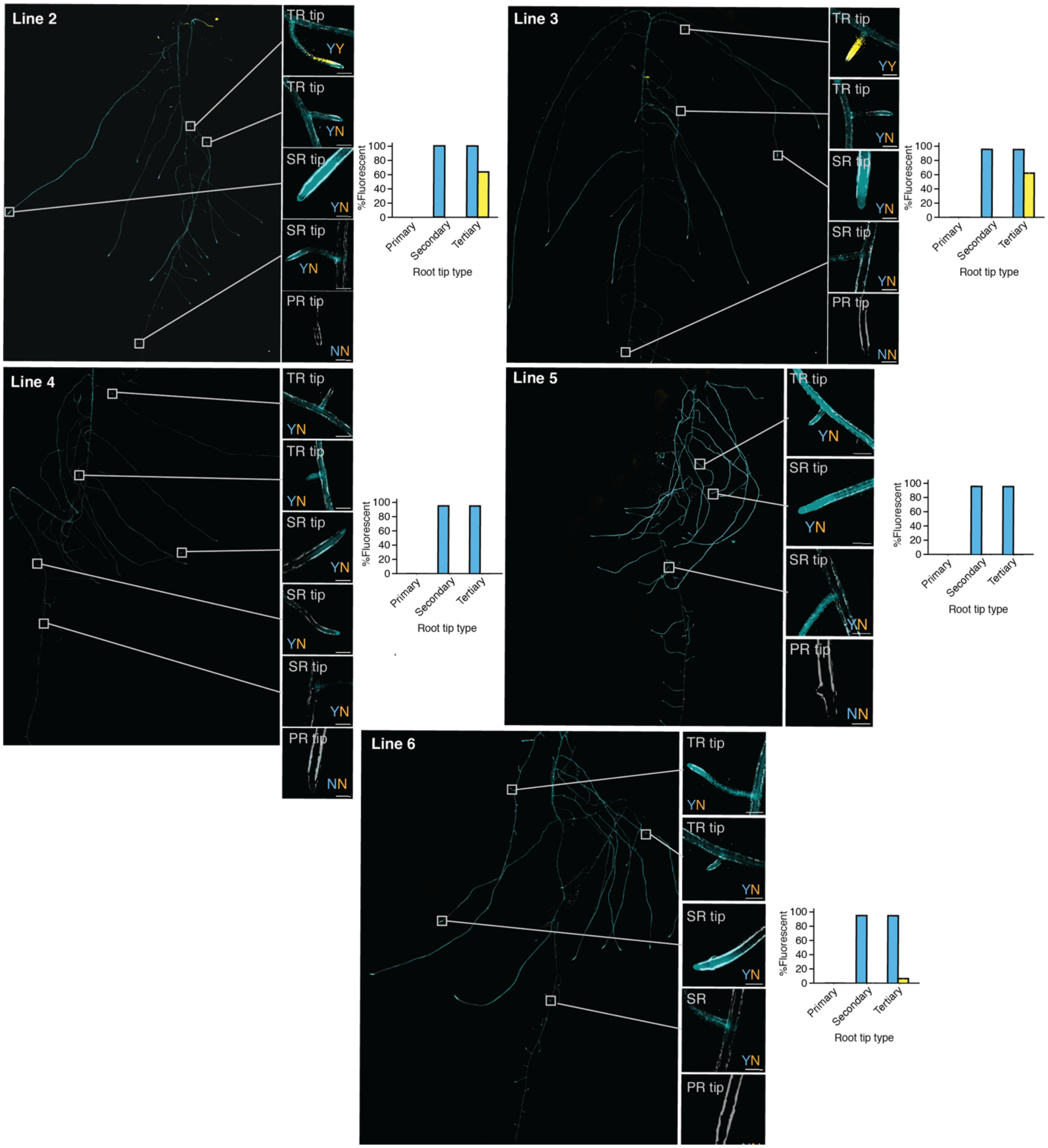


**Supplementary Figure S4. Additional T1 transgenic lines for version 3.1 counter.** Root-system fluorescence images and percentage of primary, secondary and tertiary root tips with detectable CFP or YFP signal for six additional T1 plants (Lines 2–6) carrying the version 3.1 counter (pSC096); Line 1 is shown in Supp. Fig. S3. Images show merged CFP (cyan), YFP (yellow), and reflected-light (gray) channels. Insets show secondary and tertiary root tip signal with Y and N indicating root tips scored as either fluorescence-positive (Y) or fluorescence-negative (N) for CFP (blue) and YFP (yellow). Each plot shows percentage data for one T1 plant. Signal-positive and total root tip counts are provided in Supp. Table S2, and across-plant statistical comparisons are reported in Supp. Table S3. Scale bars, 250 μm.

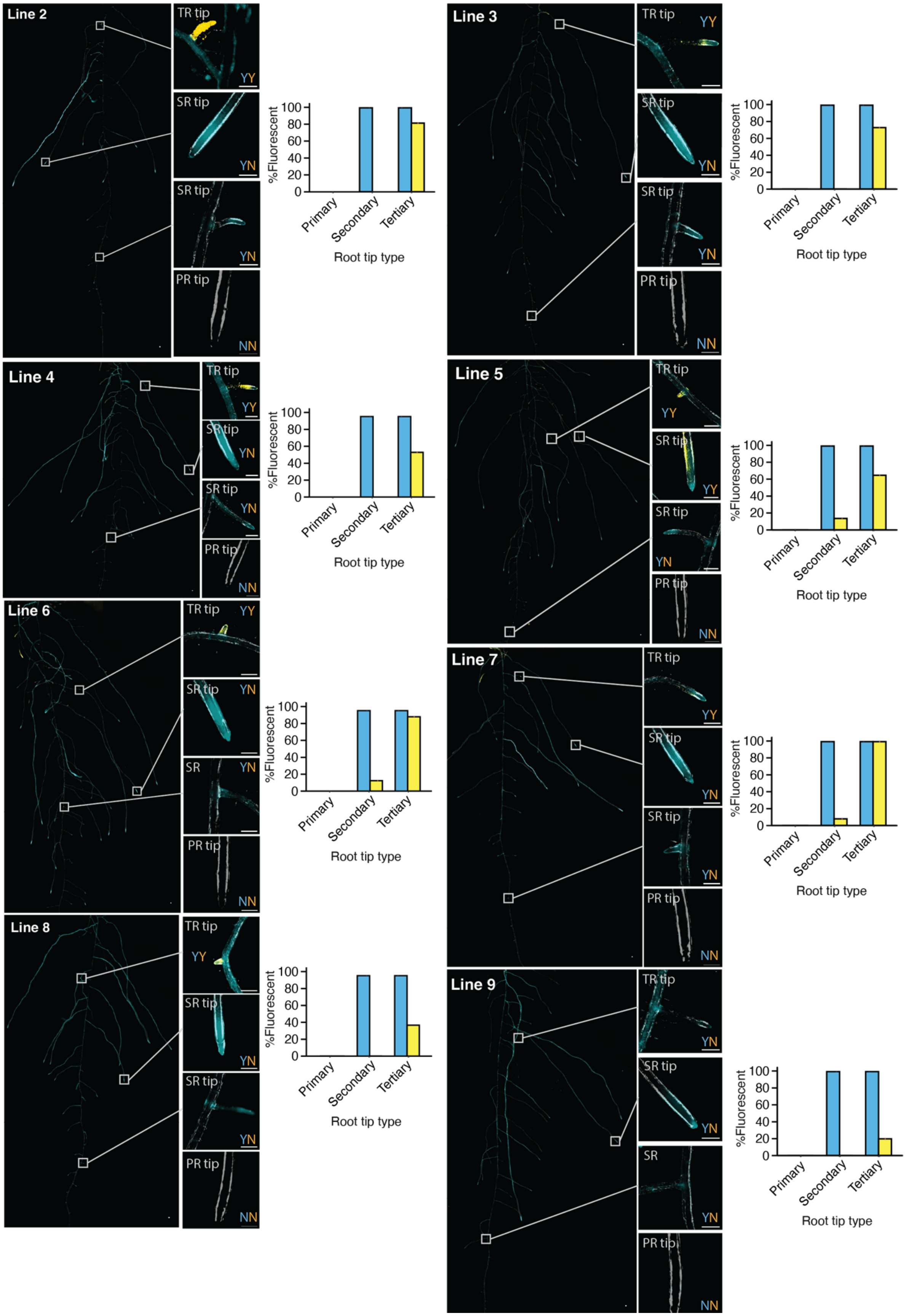

Line 2
TR tip
YY
SR tip
YN
SR tip
YN
PR tip
NN
%Fluorescent
100
80
60
40
20
0
Primary
Secondary
Tertiary
Root tip type
Line 3
TR tip
YY
SR tip
YN
SR tip
YN
PR tip
NN
Line 4
Line 5
Line 6
SR
Line 7
Line 8
Line 9
TR tip
YN

**Fig. S5. Additional version 3.2 transgenic plants and root-order fluorescence profiles.** Root-system fluorescence images and percentage of primary, secondary and tertiary root tips with detectable CFP or YFP signal for eight additional T1 plants (Lines 2–9) carrying the version 3.2 counter (pSC198); Line 1 is shown in Fig. 3. Images show merged CFP (cyan), YFP (yellow), and reflected-light (gray) channels. Insets show secondary and tertiary root tip signal with Y and N indicating root tips scored as either fluorescence-positive (Y) or fluorescence-negative (N) for CFP (blue) and YFP (yellow). Each plot shows percentage data for one T1 plant. Signal-positive and total root tip counts are provided in Supp. Table S2, and across-plant statistical comparisons are reported in Supp.Table S3. Scale bars, 250 μm.

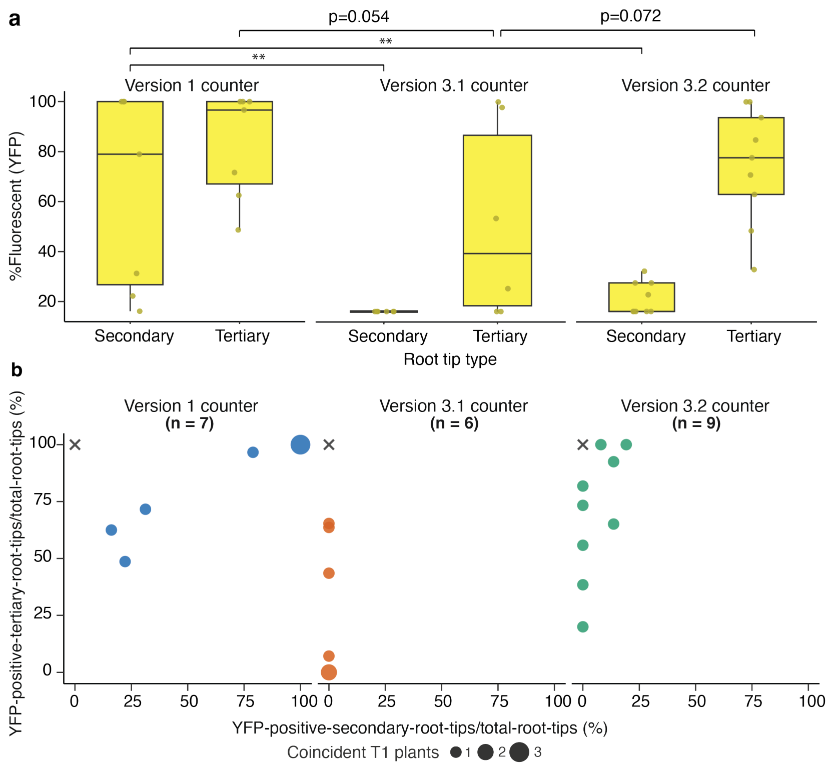


**Supplementary Figure S6. Counter trade-offs between premature secondary root flipping and tertiary root YFP output.** (a) Percent YFP-positive root tips for independent T1 transgenic plants containing the version 1 (n = 7), version 3.1 (n = 6), or version 3.2 (n = 9) counting circuit. Boxes span the first to third quartiles, central lines indicate medians, and whiskers extend to the most extreme observations within 1.5 times the interquartile range of the box limits. Each individual data point represents a single plant. Statistical significance assessed with two-sided exact Wilcoxon–Mann–Whitney rank-sum comparisons of independent construct groups, with Holm correction across all six between-version comparisons to control family-wise error. Double asterisks indicate adjusted $P < 0.01$. All six comparisons are reported in Supp. Table S3.
(b) Percent YFP-positive secondary vs tertiary root tips in each plant. Crosses mark the design target of 0% secondary-root and 100% tertiary-root YFP, not a measured observation. Counts and statistical results are provided in Supplementary Tables S2 and S3; the raw counts and analysis and plotting code are provided in Supplementary Data S5.

**Supplementary Data S1.** Raw FASTQ files for pSC178 first-transition lines analyzed in Figure 1.

**Supplementary Data S2.** Raw FASTQ files for verson1 counter (pSC235) lines analyzed in Figure 2.

**Supplementary Data S3.** Raw FASTQ files for for versions 3.1 (pSC096) and 3.2 (pSC198) lines analyzed in Figure 3 and Supplementary Figure S3.

**Supplementary Data S4.** Amplicon sequencing analysis script.

**Supplementary Data S5.** R analysis and plotting script.

**Supplementary Data S6.** micropub source-code repository.

**Supplementary Data S7.** micropub render-manifest and preset JSON files used for manuscript figures
